# Comparison of wholemount dissection methods for neuronal subtype marker expression in the mouse myenteric plexus

**DOI:** 10.1101/2023.01.17.524014

**Authors:** Julieta Gomez-Frittelli, Ryan Hamnett, Julia A. Kaltschmidt

## Abstract

**Background:** Accurately reporting the identity and representation of enteric nervous system (ENS) neuronal subtypes along the length of the gastrointestinal (GI) tract is critical to advancing our understanding of ENS control of GI tract function. Reports of varying proportions of subtype marker expression have employed different dissection techniques to achieve wholemount muscularis preparations of myenteric plexus. In this study we asked whether differences in GI dissection methods could introduce variability into the quantification of marker expression.

**Methods:** We compared three commonly used methods of ENS wholemount dissection: two flat-sheet preparations that differed in the order of microdissection and fixation as well as a rod-mounted peeling technique. We assessed marker expression using immunohistochemistry, genetic reporter lines, confocal microscopy, and automated image analysis.

**Key Results and Conclusions:** We found no significant differences between the two flat-sheet preparation methods in the expression of calretinin, neuronal nitric oxide synthase (nNOS), or somatostatin (SST) in ileum myenteric plexus. However, the rod-mounted peeling method resulted in decreased marker labeling for both calretinin and nNOS. This method also resulted in decreased transgenic reporter fluorescent protein (tdTomato) for substance P in ileum and choline acetyltransferase (ChAT) in both ileum and distal colon. These results suggest that labeling among some markers, both native protein and transgenic fluorescent reporters, is decreased by the rod-mounted mechanical method of peeling, demonstrating a critical variability in wholemount muscularis dissection methods.

## Introduction

The enteric nervous system (ENS) is an autonomous network of neurons residing in two mesh-like layers (plexuses) within the wall of the gastrointestinal (GI) tract. The submucosal plexus (SMP) lies just below the mucosa, regulating secretions and local blood flow, while the myenteric plexus (MP) resides between the circular (CM) and longitudinal (LM) muscle layers, controlling gut motility patterns^1^. Distinct regions of the GI tract perform different functions, and we are beginning to understand how this is reflected in motility patterns and the structure of the ENS controlling them^2^, especially by the involvement of many enteric neuronal subtypes^3–5^. ENS subtypes have been previously defined by their morphology^6^, electrophysiology^7^, and marker expression^3^. Recently, multiple single cell RNA sequencing (scRNAseq) studies have also defined enteric neuronal subtypes by RNA expression^4,5,8,9^. Associating these RNA signatures with previously defined ENS? subtypes is of pressing interest to the field.

Comparing results between many of the studies reported on ENS subtypes can be difficult, however. Reports of the proportion of specific subtypes or marker expression in the ENS have sometimes varied, such as reports of 6-23% of small intestine myenteric neurons expressing enkephalin^3,10^. Variability such as this can result from animal model^3^, background^11^, age^12^, or GI tract region analyzed^13^.

Tissue processing has also been shown to introduce variability in the quality of enteric marker visualization^14^. Many different GI dissection methods and adaptations thereof have been reported over the years, especially for wholemount preparations of the myenteric plexus (*see Table 1*). Here, we assessed this potential source of variability in myenteric neuron marker expression in a controlled set of experiments testing (i) the order of microdissection and fixation and (ii) the usage of a rod during the microdissection (see *Detailed dissection protocols*) across two landmark-defined intestinal regions, ileum and distal colon. We find no significant differences in labeling between two flat sheet methods differing in the order of dissection and fixation steps, but report significantly decreased labeling using the rod peeling method across most markers and regions tested.

**Table 1:** Wholemount muscularis dissection methods and variations.

| Method | Variations |
| --- | --- |
| Fix segments pinned flat, then peel muscularis<sup>13</sup> | Muscle relaxants (nicardipine)<sup>6,23,24</sup> |
|  | Amount of stretch<sup>23,25</sup> |
|  | Layers peeled<sup>3,26</sup> |
|  | Transcardial perfusion with fixative<sup>27</sup> |
|  | Microdissection vs razor blade<sup>27</sup> |
| Peel muscularis, then fix<sup>28</sup> | Layers peeled<sup>2,29</sup> |
|  | No fixation (for fluorescent reporters)<sup>22</sup> |
|  | Transcardial perfusion with fixative<sup>22</sup> |
| Peel on rod, then fix<sup>30</sup> | Peeling longitudinal muscle – myenteric plexus (LMMP) only (in rats)<sup>31</sup> |
| Fix as tube, then cut open and peel | Distend gut with Krebs<sup>32</sup> or fixative<sup>32</sup> |
|  | No distension<sup>33</sup> |

## Materials and Methods

### Mice

All procedures conformed to the National Institutes of Health Guidelines for the Care and Use of Laboratory Animals and were approved by the Stanford University Administrative Panel on Laboratory Animal Care. Mice were group housed up to a maximum of five adults per cage. Food and water were provided *ad libitum* and mice were maintained on a 12:12 LD cycle.

Experiments used wild-type C57BL/6J mice and the reporter lines choline acetyltransferase(ChAT)-IRES-Cre (Δneo)^15^ and Tachykinin precursor 1(Tac1)-IRES2-Cre-Δ^16^, both crossed to Cre-dependent tdTomato (tdT) Ai14 mice^17^ (hereafter ChAT-Cre-tdT and Tac1-Cre-tdT, respectively). Mice aged 2-4 months from both sexes were used, except for ChAT-Cre-tdT (only males) and Tac1-Cre-tdT (only females).

### Dissection protocols

#### Intestinal segment harvest

Retrieval of Ileum and distal colon segments was performed as in (Hamnett, Dershowitz *et al*., 2022)^13^ to the point of removing the mesentery from the intestines. The dissection then proceeded with *Method 1, Method 2,* or *Method 3*, described below.

#### Method 1: Fix flat then peel

The intestinal segment was opened along the mesenteric border, flipped over (serosa-up) and pinned flat along the edges in a Sylgard-bottomed glass Petri dish, stretched under light tension. Segments were fixed in 4% PFA at 4°C for 90 minutes with gentle rocking, followed by three washes with cold PBS for at least 10 minutes each. To obtain muscularis with myenteric plexus, the muscularis was carefully peeled up and away from the submucosa/mucosa, separating the layers along the first 2-3 mm. The segment was flipped over, mucosa side up, and re-pinned. The mucosa/submucosa was carefully pulled away, while a cotton-tipped swab was used to gently hold down the muscularis. Segments were stored in PBS with 0.1% NaN3 at 4°C until use.

#### Method 2: Peel flat then fix

Each segment was prepared as in *Method 1* with the following modifications: (i) the order of microdissection and fixation of the pinned segments was reversed and (ii) fine forceps were used to grasp the mucosa/submucosa and gently pull it away, avoiding the use of the cotton swab applying pressure to the muscularis before fixing. Muscularis segments were fixed, washed, and stored as in *Method 1*.

#### Method 3: Peel on rod

A glass rod was inserted through the lumen of the full-length intestinal segment. The diameter of the glass rod (2 mm) was chosen to fit the diameter of the segment and slightly distend it. Forceps were used to gently perforate the muscularis (but not the mucosa/submucosa) at the mesenteric attachment. Ice-cold PBS was applied frequently during dissection to maintain moisture and temperature of the segment. A cotton-tipped swab moistened with cold PBS was used to gently peel away the muscularis, swabbing circumferentially from the perforated mesenteric edge while the segment was manually held stable on the rod. Fully peeled muscularis segments were transferred to a Sylgard-bottomed glass Petri dish with cold PBS and pinned flat along the edges under light tension. Muscularis segments were fixed, washed, and stored as in *Method 1*.

### Immunohistochemistry

Immunohistochemistry was performed as described previously^13^. All wash steps were at least 10 minutes each. The following antibodies were used: rabbit anti-calretinin (1:4000; Millipore AB5054), goat anti-calretinin (1:8000; Swant SG1), rat anti-SST (1:500; Millipore MAB354), rabbit anti-nNOS (1:1000; Sigma N7280), and human anti-ANNA1 (HuC/D) (1:50,000; Mayo Clinic).

### Confocal imaging

Images were acquired as described previously^13^. Z-stacks were acquired with 2.5-3 μm between each focal plane.

### Neuronal and marker quantification

Image analysis was performed using ImageJ/FIJI (NIH, Bethesda, MD), as described previously^13^. SST was counted manually after being combined with the thresholded HuC/D image stack and identified by its distinctive cytoplasmic expression pattern.

### Statistical analyses

Statistical tests and graphical representation of data were performed using Prism 9 software (GraphPad). Statistical comparisons were performed using one-way ANOVA followed by Tukey’s correction for multiple comparisons to assess if variations in dissection method were a significant factor (*p*<0.05) for calretinin, nNOS, and SST subtype marker proportion. Two-way ANOVAs followed by Šídák’s multiple comparisons tests were used to determine the effect of dissection method on ChAT-Cre-tdT and Tac1 - Cre-tdT marker subtype proportion. Asterisks indicate significant differences.

## Results

To probe possible effects of different tissue dissection methods on marker expression, we separately tested significantly different aspects of three dissection methods described in the field (detailed in *Methods*). We focused on (i) the order, chronologically, of peeling and fixation, and (ii) mechanically different methods of peeling the muscularis, while preserving variables such as dissection solution and temperature, type of fixative and duration of fixation across the three methods.

We first analyzed neuronal marker expression via immunohistochemistry. We chose three subtypes of enteric neurons which can be identified by expression of distinct marker proteins for which antibody staining yields easily quantifiable cell body labeling: excitatory motor neurons by calretinin expression, inhibitory motor neurons by neuronal nitric oxide synthase (nNOS) expression, and a subpopulation of interneurons by somatostatin (SST) expression^18–20^. For all three of these markers, no significant difference in proportion of neurons labeled was found between the two flat-sheet microdissection methods (*Methods 1 and 2*) (Fig 1). In the case of SST, there was also no significant difference in proportion of neurons labeled using the rod peeling method (*Method 3*) compared to the flat-sheet methods (Fig 1l). However, for both calretinin (Fig 1d) and nNOS (Fig 1h), the rod peeling method resulted in significantly fewer neurons labeled than either flat sheet method (calretinin, ~35% (*Methods 1 and 2*) decreased to ~10% (*Method 3*); nNOS, ~23% (*Methods 1 and 2*) decreased to ~13% (*Method 3*)).

**Figure 1.**
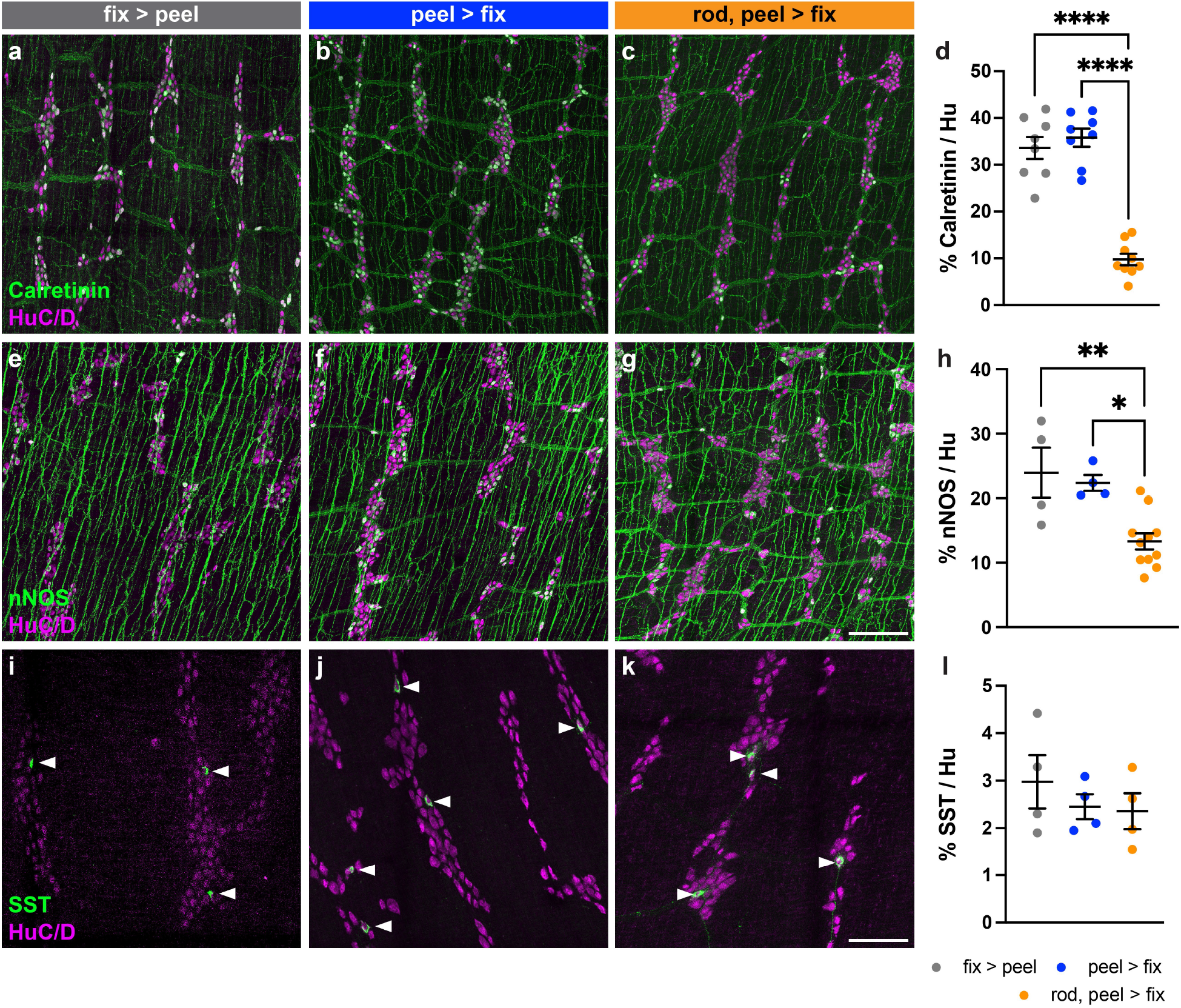
Proportion of ileum myenteric plexus neurons labeled immunohistochemically by three dissection methods. (**a-c**) Representative images of ileum myenteric plexus wholemounts prepared via flat-sheet fixing then peeling (fix > peel) (**a**), flat-sheet peeling then fixing (peel > fix) (**b**), or rod peeling then fixing (rod, peel > fix) (**c**), immunohistochemically labeled for calretinin (green) and neuronal label HuC/D (magenta). (**e-g**) As in (**a-c**), labeled for nNOS (green). (**i-k**) As in (**a-c**), labeled for SST (green). (**d, h, l**) Proportion of total HuC/D neurons (mean ± SEM) positive for each neuronal marker for each dissection method. (**d**) Calretinin: fix > peel, n = 8; peel > fix, n = 8; rod, peel > fix, n = 9. (**h**) nNOS: fix > peel, n = 4; peel > fix, n = 4; rod, peel > fix, n = 11. (**l**) SST: fix > peel, n = 4; peel > fix, n = 4; rod, peel > fix, n = 4. Scale bar represents 200 μm in (**a-c**), (**e-g**); 100 μm in (**i-k**). All tests one-way ANOVA. \**p*<0.05, \*\**p*<0.01, \*\*\*\**p*<0.0001.

We next asked whether the mechanical technique of wholemount peeling would also impact the reporting of genetically labeled neurons in transgenic mice. Transgenic mice that express a reporter fluorescent protein in a gene-dependent manner can be used to quantify neuronal subtype marker expression for markers where antibody staining does not yield easily quantifiable cell body labeling, such as ChAT and substance P. Like calretinin, ChAT and substance P are both markers for excitatory motor neurons. We compared the two mechanically different peeling methods, flat-sheet (*Method 1*) and rod mounted (*Method 3*) in ChAT-Cre-tdT and Tac1-Cre-tdT mice, and expanded our analysis to two regions, ileum and distal colon. In Tac1-Cre-tdT ileum the proportion of neurons labeled by tdT was similar between the two methods tested (Fig 2 f-g, j); however, in ChAT-Cre-tdT ileum, the proportion of neurons labeled was significantly decreased with the rod peeling method as compared to the flat-sheet method, from ~62% (*Method 1*) to ~26% (*Method 3*) (Fig 2 a-b, e). In the distal colon, the rod peeling method resulted in decreased tdT labeling in ChAT-Cre-tdT (~38% (*Method 1*) decreased to ~14% (*Method 3)*) and Tac1-Cre-tdT (~36% (*Method 1*) decreased to ~3% (*Method 3)*) myenteric plexus, with labeling in Tac1-Cre-tdT colon almost completely abolished (Fig 2 c-e, h-j). Importantly, we did not note any differences in overall neuronal density, their general soma morphology revealed by HuC/D staining, or the subcellular localization of HuC/D protein (Fig 2 k-o), which if found might have suggested neuronal cell death or mechanical damage^21^ as an underlying cause for decreased labeling. Taken together, our results demonstrate that while the order of dissection and fixation steps in flat-sheet methods does not impact marker labeling, for both native protein detection and transgenic reporter fluorescent protein expression, and in multiple intestinal regions, the rod peeling method can result in decreased marker labeling compared to the flat-sheet microdissection methods.

**Figure 2.**
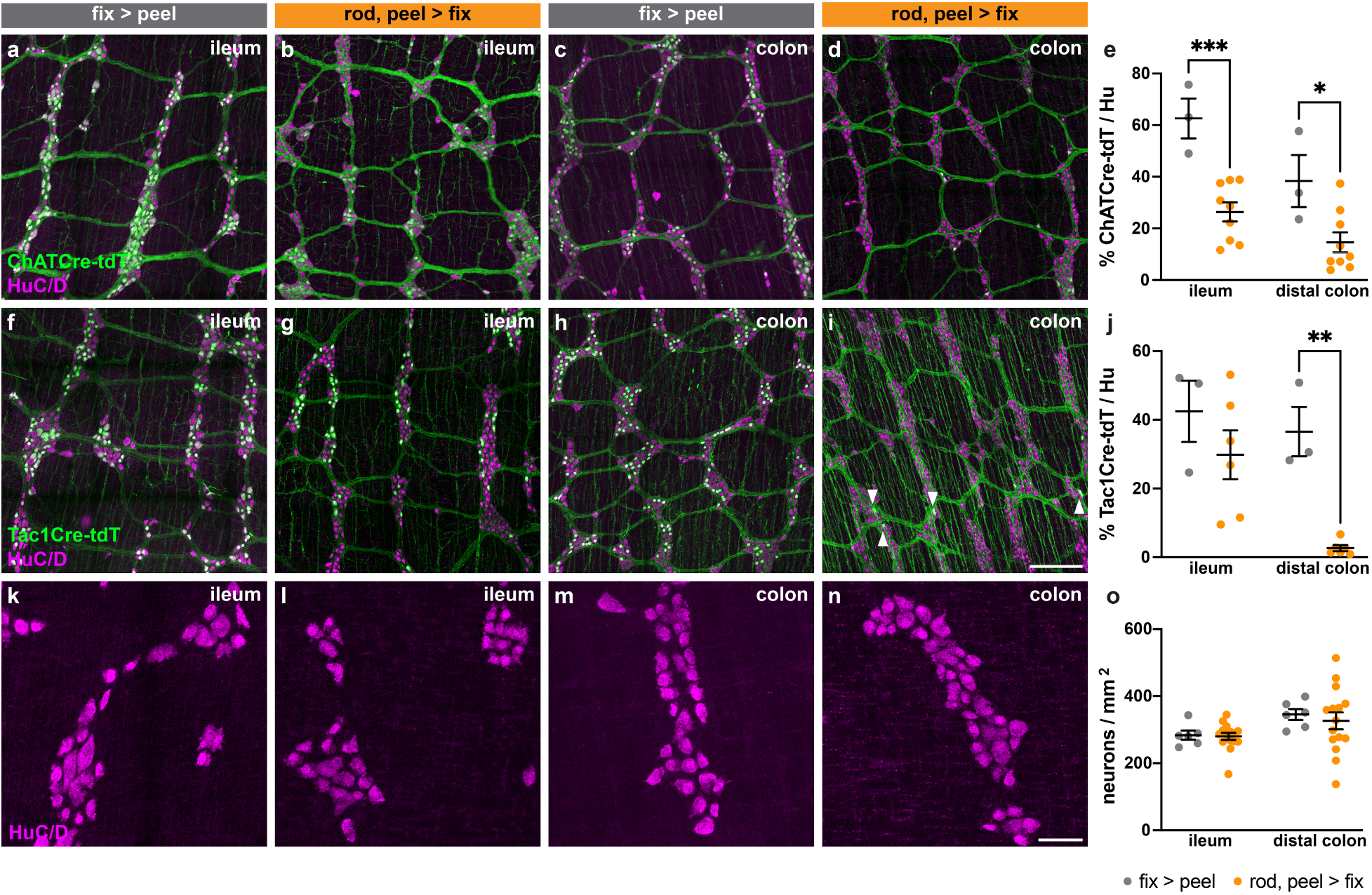
Proportion of ileum and distal colon myenteric plexus neurons labeled genetically by two distinct dissection methods. (**a-d**) Representative images of ileum (**a, b**) and distal colon (**c, d**) myenteric plexus prepared via flat-sheet fixing then peeling (fix > peel) (**a, c**), or rod peeling then fixing (rod, peel > fix) (**b, d**), genetically labeled for ChAT-Cre-tdT (green) and neuronal label HuC/D (magenta). (**f-i**) As in (**a-d**), labeled for Tac1-Cre-tdT (green). (**e, j**) Proportion of total HuC/D neurons (mean ± SEM) positive for each neuronal marker for each region and each dissection method. (**e**) ChAT-Cre-tdT: ileum fix > peel, n = 3; ileum rod, peel > fix, n = 9; distal colon fix > peel, n = 3; distal colon rod, peel > fix, n = 9. (**j**) Tac1-Cre-tdT: ileum fix > peel n = 3; ileum rod, peel > fix n = 6; distal colon fix > peel n = 3; distal colon rod, peel > fix n = 6. (**k-n**) As in (**a-d**), showing HuC/D only. (**o**) Density of HuC/D neurons (mean ± SEM) for each region by each dissection method. All tests two-way ANOVA. Scale bar represents 200 μm in (**a-d**), (**f-i**); 50 μm in (**k-n**). All tests two-way ANOVA. \**p*<0.05, \*\**p*<0.01, \*\*\**p*<0.001.

## Discussion

In this study we performed a controlled and comparative investigation of the effect of different wholemount mouse myenteric plexus preparations on neuronal subtype marker expression as a proportion of total neurons. We evaluated whether marker expression is dependent on either (i) the chronological order of the dissection and fixation steps, or (ii) the mechanical method of wholemount dissection (peeling from flat tissue or peeling on a rod). We found no significant differences in neuronal labeling when varying the order of the peeling and fixation steps in flat-sheet preparations. By contrast, in our hands, we found that some neuronal markers (both native proteins and transgenic reporter proteins) are affected by the mechanical method of wholemount peeling, and significantly decreased in the rod peeling method. This was particularly striking in the case of ChAT-Cre-tdT, where the proportion of neurons labeled was decreased from about two-thirds (~62%) to about one-quarter (~26%) in ileum myenteric plexus. ChAT and nNOS are known to cover the vast majority of enteric neurons between them^22^, but the decrease in both ChAT-Cre-tdT and nNOS labeling with the rod peeling method would suggest a large proportion of myenteric neurons unlabeled for either.

The exact mechanism of the described differences in marker labeling is unclear. There may be some as yet unknown effect introduced by the distension of the intestinal segment on the glass rod or the handling of the muscularis as it is peeled away from the mucosa, which may cause a rapid change in neuronal marker protein expression without obvious evidence of cell death or mechanical damage. In this study we did not explicitly test for any effect of time elapsed before fixation; however, in our hands the time to fixation between the flat-sheet peel pre-fixation method and the rod peeling method was comparable, suggesting that any decrease in marker labeling with the rod peeling method is not due to the time required to process the tissue prior to fixation. Finally, we note that our comparative study is not exhaustive and that future experiments are required to uncover the cause underlying the observed differences in marker expression.

## Acknowledgments

We thank the members of the Kaltschmidt lab for experimental advice and discussions. We thank Vanda Lennon (Mayo Clinic) for the HuC/D primary antibody, and William Giardino for providing the Tac1-Cre mouse. This work was supported by the Stanford ChEM-H Chemistry/Biology Interface Predoctoral Training Program and the National Institute of General Medical Sciences of the National Institutes of Health Award T32GM120007 (J.G.F.), an EMBO Fellowship ALTF 180-2019 (R.H.), and the Wu Tsai Neurosciences Institute, the Stanford University Department of Neurosurgery and research grants from The Shurl and Kay Curci Foundation, The Firmenich Foundation and The Carol and Eugene Ludwig Family Foundation (J.A.K.).

